# Rac1 Selectively Binds a Specific Lamellipodin Isoform via a Noncanonical Helical Interface

**DOI:** 10.1101/2025.06.24.661328

**Authors:** Tong Gao, Pingfeng Zhang, Alison M. Kurimchak, James S. Duncan, Jinhua Wu

**Affiliations:** Cancer Signaling and Microenvironment Program, Fox Chase Cancer Center, Philadelphia, PA 19111, USA

## Abstract

Lamellipodin (Lpd) is a multifunctional adapter protein that regulates cell migration and adhesion by recruiting Ena/VASP proteins to the leading edge and modulating actin polymerization. The interaction of Lpd and Rho family or Ras family GTPases is crucial for regulating actin dynamics. Contrary to previous assumptions that the main Lpd isoform interacts with Rac1, here we show that strong and specific binding to Rac1 is instead mediated by the short isoform Lpds. This interaction is dependent on Rac1’s GTPase activity and a short insertion (cs2) within the coiled-coil (CC) region unique to the Lpds isoform. Structural modeling and mutagenesis analyses further reveal that Lpds engages Rac1 through a noncanonical, single-helix binding mode distinct from the classical helical pair configuration. Our results reveal a novel isoform-dependent GTPase:effector binding mode and suggest a critical regulatory pathway that may represent a promising therapeutic target in Rac1-driven cancer progression.

## Introduction

Cell migration and adhesion are fundamental biological processes essential for diverse functions including immune responses, tissue development, cell proliferation, and cancer progression. These processes are mainly controlled by actin polymerization and cytoskeleton remodeling at the leading edge of migrating cells in lamellipodia and filopodia. Disruption or misregulation of these processes are linked to pathological conditions including tumor metastasis and invasion. Among key actin-binding proteins, the Ena/VASP (enabled/vasodilator-stimulated phosphoprotein) family proteins play important roles in regulating actin dynamics. The Ena/VASP proteins are recruited to the leading edge of cells by lamellipodin (Lpd), a member of the MRL (Mig10/RIAM/Lpd) family of adaptor proteins.^1,2^

The MRL proteins are multi-adaptor proteins that share similar domain architectures but serve diverse roles in regulating cell migration and cell adhesion. In mammals, the two MRL ortholog proteins, Lpd and Rap1-GTP-interacting adaptor molecule (RIAM), both contain a conserved center domain composed of a Ras-associating (RA) and a pleckstrin homology (PH) structural modules, known as the RA-PH domain. They also possess an N-terminal talin-binding sites, a coiled-coil region preceeding the RA-PH domain, which is followed by several polyproline motifs responsible for interacting with profiin and Ena/VASP.^3^ The RA-PH domain, found in MRL and Grb family proteins, interacts with small GTPase and phosphatidylinositol 4,5-bisphosphate (PIP_2_) to regulate cell migration and protrusion formation.^3,4,5,6^ Both Lpd and RIAM play crucial roles in integrin activation by recruiting talin to the plasma membrane (PM), effectively linking small GTPase signaling to essential cellular processes such as cell adhesion. ^5,7^

Small GTPases are a family of guanine nucleotide-binding proteins that function as molecular switches to regulate diverse cellular processes in eukaryotes, including cell proliferation, differentiation, and cytoskeletal dynamics. Based on their sequence homology, structure, and function, they can be divided into five major families, Ras, Rho, Ran, Rab, and Arf.^8^ Among them, Ras GTPases are well-known oncoproteins implicated in tumorigenesis. Rho GTPases, including Rac1, RhoA, and Cdc42, are recognized for their role in regulating cell cycle, cell polarity and cell migration and play key roles in cancer metastasis. ^9–12^ As a prototype member of the Rho family, Rac1 regulates cytoskeleton reorganization and lamellipodia formation. Elevated Rac1 activity is linked to enhanced metastasis and tumorigenesis (PMC6261428). Recent studies have indicated Rac1 as a potential therapeutic target due to its important role of Rac1 in driving metastasis and resistance to therapies. ^13–15^

Lpd has been shown to interact with two small GTPases, M-Ras in the Ras family and Rac1 in the Rho family.^16,17^ In cortical neurons, Lpd directly binds to M-Ras and mediates dendritic growth and branching of neuronal dendrite. This process is regulated by Plexin through its GTPase-activating protein (GAP) activity.^18^ As a Rac1 GTPase effector. Lpd localizes to the leading edge of migrating cells, specifically at the tips of lamellipodia and filopodia, where it recruits Ena/VASP protein to the growing protrusions, thus enhancing cell migration by accelerating actin polymerization.^1,19,20^ Moreover, Lpd also interacts with other actin-regulatory proteins. In mouse embryonic fibroblast, Lpd interacts with Scar/WAVE (Suppressor of cAMP Receptor/WASP-family Verprolin-homologous) complex, a key Rac1 effector that promotes Arp2/3-mediated actin branching. This interaction is positively regulated by activated Rac1 and is essential for lamellipodial extension and cell migration. ^21^

Many studies have indicated that Lpd is linked to various cancer-related processes. Lpd has been shown to promote glioblastoma invasion, proliferation and radiosensitivity through epidermal growth factor receptor (EGFR) signaling axis.^22^ In breast cancer, Lpd enhances cell migration by regulating Ena/VASP proteins and the Scar/WAVE complex. ^2^ In oral squamous cell carcinoma (OSCC) cell lines, Lpd expression is upregulated upon cell migration induced by Hepatocyte growth factor (HGF)/c-Met signaling. ^23^ Moreover, in osteosarcoma, reduced expression of Lpd results in tumor suppression.^24^ All those studies demonstrate that Lpd not only functions in maintaining normal actin dynamics and cell adhesion but also plays important roles in cancer progression and metastasis.

Previous studies have shown that CED-10, the *C. elegans* ortholog of Rac1, interacts strongly with MIG-10, the ortholog of Lpd.^25^ However, direct interaction of mammalian Lpd or its RA-PH with Rac1 has not been explicitly demonstrated. Interestingly, our data suggested that neither full length Lpd nor its isolated RA-PH domain exhibits significant binding to activated Rac1. In this study, we identified a specific Lpd isoform, known as Lpds, that displays remarkably high binding affinity for active Rac1.

Through domain-mapping and mutation studies, we demonstrated that a short splicing insertion near the coiled-coil region of Lpds is required for this high-affinity interaction. While the binding is Rac1 activity-dependent, our biochemical data and structural analyses suggest that the Lpds interacts with Rac1 through a previously uncharacterized interaction, rather than the canonical intermolecular β-sheet interaction for typical RA-GTPase interaction or the helical pair interaction seen in some Rho family GTPases and effectors.^26^ Our results reveal an isoform-specific GTPase effector signaling mechanism, and propose a novel GTPase-effector interacting architecture.

## Results

### Activated Rac1 specifically interacts with one of Lpd isoforms

Although the interaction of *C. elegant* orthologs of Lpd and Rac1 has been demonstrated,^25^ no significant interaction of mammalian Lpd and Rac1 was detected in our preliminary data. Lpd possesses several isoforms due to alternative splicing.^19^ The splicing variants often share many conserved functional regions, with some containing additional segments.^27^ To date, at least nine Lpd isoforms have been identified in human. The main Lpd isoform consists of 1250 amino acids, including an N-terminal talin binding site (TBS), an autoinhibitory segment (IN), a coiled-coil (CC) region, the RA-PH domain, and an unstructured C-terminal ENA-VASP binding tail (**Fig. 1A**).^19,28,29^ The other eight isoforms lack the C-terminal ENA-VASP binding tail. In isoforms Q70E73-3, -4, -6, -7, -8, and -9, alternative RNA splicing results in two additional helical segments upstream of the CC region. We therefore refer to these insertions as coiled segments (CS), comprising cs1 and cs2.

**Figure 1.**
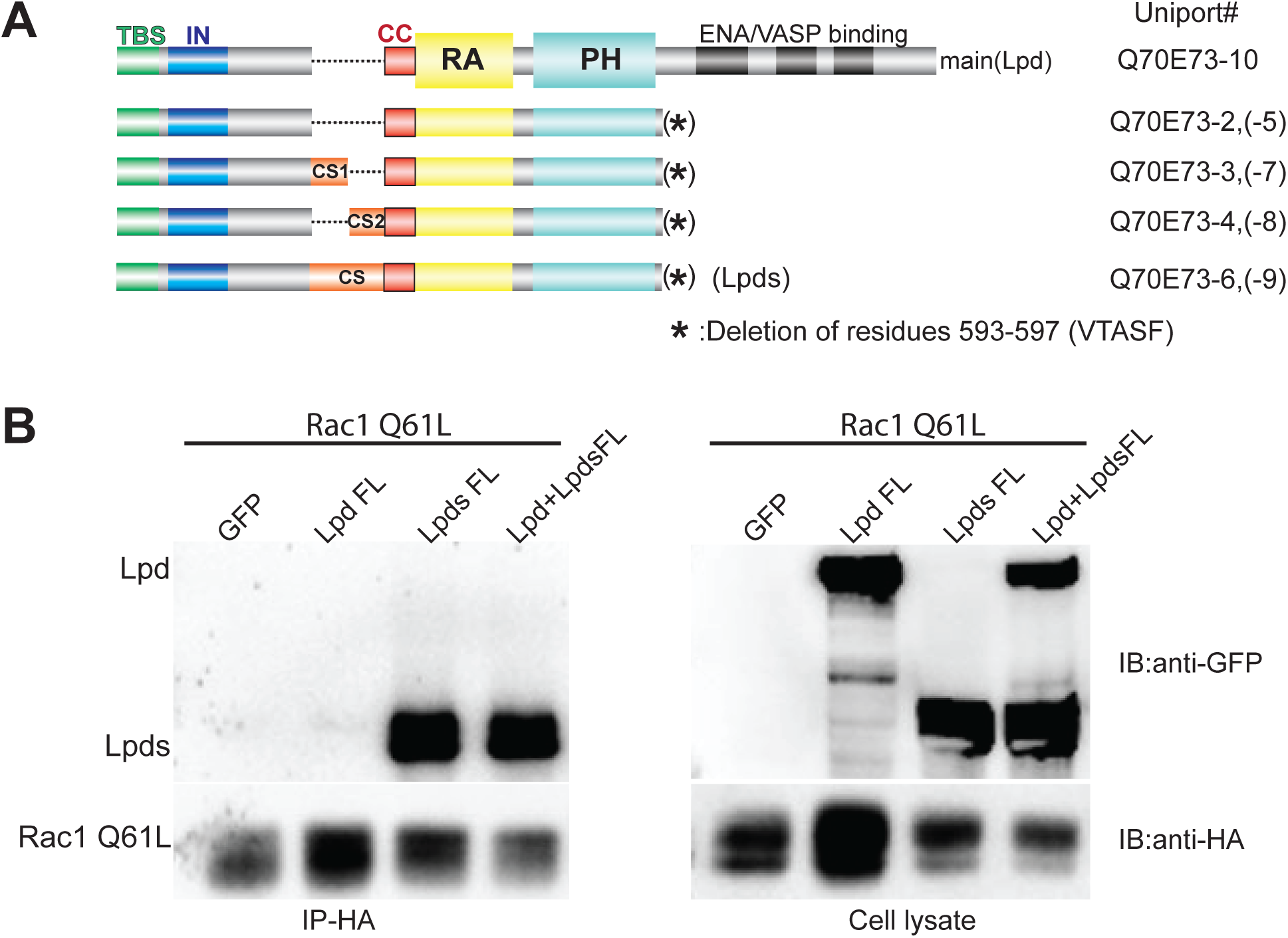
The Lpds isoform of Lpd has strong binding affinity with activated Rac1 specifically. (**A**). Schematic representation of Lpd isoforms. TBS: talin binding site; IN: autoinhibitory segment; CC: coiled-coil region; CS: insertion of cc; RA, Ras-association domain; PH, pleckstrin homology domain. A talin-binding site (green) is located at the N-terminal end of Lpd and is conserved across all isoforms. The CS is depicted in orange, the CC is in red, the RA (Ras-associating) domain is in yellow, and the PH (pleckstrin homology) domain is in cyan. (**B**). The Co-IP assay of GFP-tagged full-length Lpd and Lpds and HA-tagged constitutively active Rac1 form (Rac1 Q61L). Cell lysate and IP samples are detected by Western blot using anti-HA antibody and anti-GFP antibody.

To examine the impact of the additional CS on Rac1 interaction, we performed co-IP assays to compare the binding affinities of Rac1 with the main isoform of Lpd and a short isoform, Lpds, which contains the CS insertion. Both GFP-tagged full-length Lpd and Lpds were expressed at similar levels in HEK293 cells and subjected to co-IP with activated Rac1. While no significant interaction was detected with the main isoform, the Lpds isoform co-immunoprecipitated robustly with activated Rac1 (**Fig. 1B**). The identity of Lpds was confirmed by mass spectrometry. These findings indicate that Rac1 preferentially binds the Lpds isoform, revealing an isoform-specific interaction.

Previous studies also demonstrate that Lpd can dimerize through an anti-parallel coiled-coil interaction involving the CC region.^4^ This dimerization interaction is crucial for ENA/VASP clustering at the leading edge of protrusions thus playing an important role in mediating cell migration.^1,30^ To investigate if the dimerization through the CC region contributes to Rac1 binding, we co-expressed Lpds, Lpd, and Rac1 in HEK293 cells. If Lpd and Lpds form heterodimer, both isoforms are expected to coimmunoprecipitate with Rac1 despite the lack of direct interaction of Lpd and Rac1. Nevertheless, only Lpds, but not Lpd, was co-immunoprecipitated with Rac1 (**Fig. 1B**). The result suggests that the Lpds interact with Rac1 as a monomer, and the dimerization through the CC region does not contribute significantly to in the interaction of Lpds and Rac1.

### Activated Rac1 binding to Lpds Requires the CS insertion

The RA-PH module is seen in Grb family and MRL family protein.^4,31–33^ The RA domain typically mediates interactions with small GTPases such as the Ras or Rap GTPases. Previous studies show that both M-Ras and Rac1 can interact with Lpd.^18,21^ To investigate whether Lpds can interact with other small GTPases, we examined the binding of Lpds with small GTPases from various Ras families. Among which, M-Ras, H-Ras and R-Ras are classical members of Ras family cancer therapy; ^8,34,35^ Rap1 belongs to the Rap subfamily, which is involved in integrin activation and plays key roles in cell adhesion; ^4^ and Rac1is an important members of Rho family that regulates cytoskeleton dynamics.^9^ A Lpds construct (1-PH) containing the intact RA-PH module and the entire sequence upstream of the module was co-expressed with each of the small GTPases in HEK293 cells for co-immunoprecipitation. While Lpds and Rac1 exhibited robust interaction, the Co-IP results showed no significant interaction between Lpds and the constitutively active Rap1, M-Ras, H-Ras or R-Ras (**Fig. 2A**). Our results suggest that Lpds preferentially associates with Rac1, rather than Rap or Ras family members, in modulating specific cellular processes.

**Figure 2.**
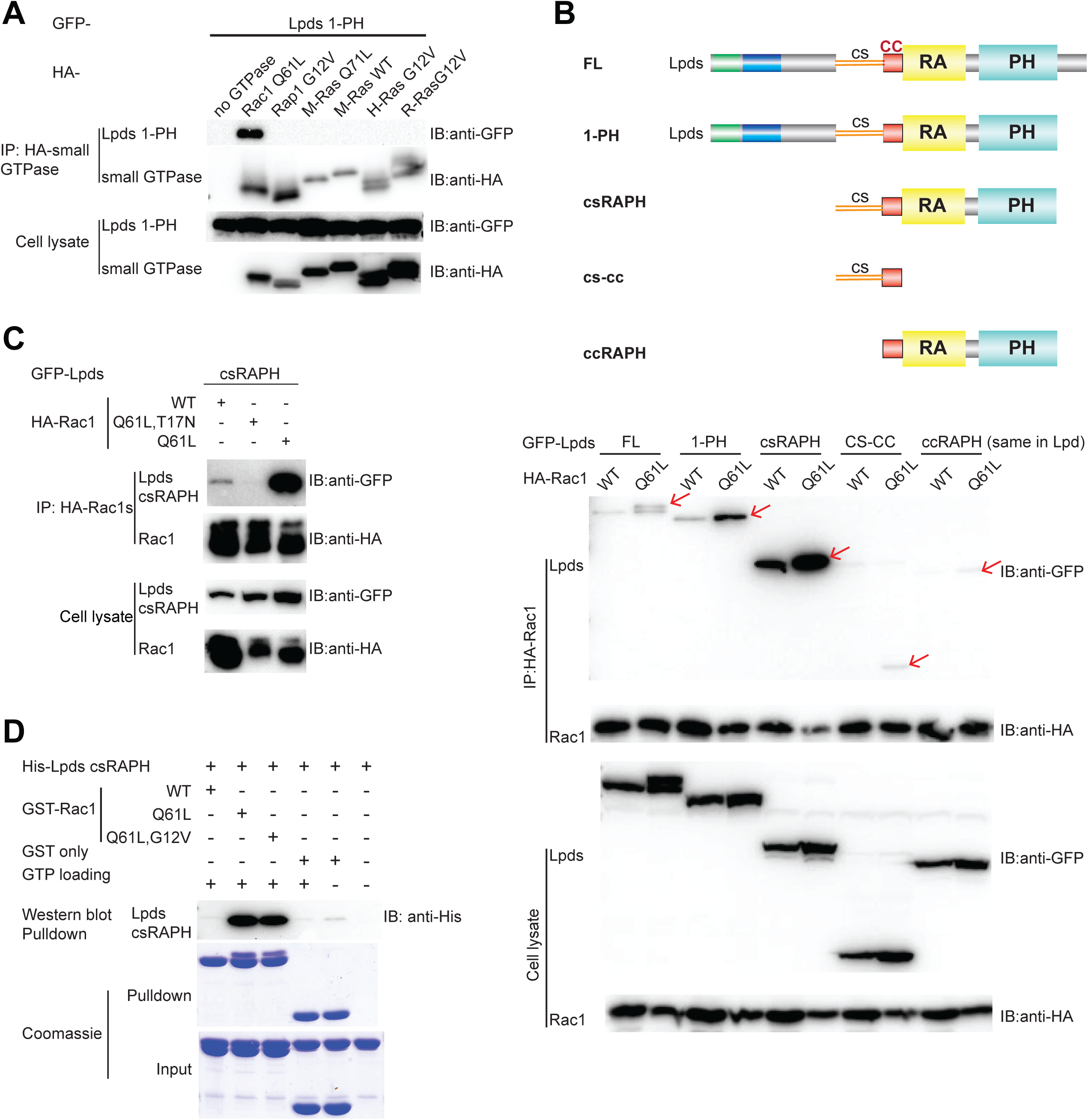
Rac1:Lpds interaction requires the GTPase activity of Rac1 and the CS insertion of Lpds. (**A**). Co-IP assay of GFP-tagged Lpds1-PH with HA-tagged Rac1 Q61L, Rap1 G12V, M-Ras Q71L, M-Ras WT, H-Ras G12V and R-Ras G12V. (**B**). Co-IP assay of HA-tagged wild-type Rac1 and Rac1 Q61L with GFP-tagged Lpds FL, 1-PH, csRAPH, cs-cc and Lpd ccRAPH. The schematic diagrams of the constructs are shown above, with the same domain color codes as in Figure 1A. (**C**). Co-IP assay of HA-tagged wild-type Rac1, constitutively actived Rac1 form (Rac1 Q61L) and inactivated Rac1 form (Rac1 Q61L/T17N) with GFP-tagged Lpds csRAPH. (**D**). Pulldown assay of GST-tagged wild-type Rac1, Rac1 Q61L and Rac1 Q61L/G12V with His-tagged Lpds csRAPH.

We further mapped the Rac1-interaction sites in Lpds by generating a serial of truncation constructs. Lpds shares the same sequence with the main Lpd isoform except for the additional CS insertion and the absence of the C-terminal ENA-VASP binding region. In addition to the full-length (FL) and 1-PH constructs of Lpds, we generated csRAPH, CS-CC, and RAPH constructs. The csRAPH construct contains only the CS, CC, and RA-PH segments; the CS-CC construct includes the CS and CC segments; and the ccRAPH construct, which is also present in the main Lpd isoform, contains the CC and RA-PH segments. (**Fig. 2B**). Co-immunoprecipitation of these constructs with wild type (WT) or constitutively active Rac1 (Q61L) showed that both the Lpds FL and Lpds 1-PH constructs interacted with Rac1. In contrast, the CS-CC or ccRAPH constructs exhibited minimal interaction. Notably, the csRAPH construct, which contains both the CS insertion and the RAPH domain, displayed the strongest binding to Rac1 among all the Lpds constructs (**Fig. 2B**). These results suggest that the CS insertion is essential for promoting Rac1 binding to Lpds and strengthening their interaction.

We next examined whether the GTPase activity of Rac1 is essential for its interaction with Lpds. We used the csRAPH construct of Lpds to test its interaction with WT, constitutively active (Q61L) and inactive (Q61/T17N) Rac1. Co-immunoprecipitation assays demonstrated that WT Rac1 binds to Lpds csRAPH, while the constitutively active Rac1 (Q61L) exhibits much enhanced binding with Lpds csRAPH. In contrast, the interaction with Lpds is abolished with the inactive Rac1 mutatnt (Q61L/T17N) (**Fig. 2C**). To rule out the possibility of indirect or mediator-assisted interactions through other cellular components, we purified His-tagged csRAPH and GST-tagged Rac1 proteins and performed *in vitro* pulldown assays. GTP-loaded, constitutively active Rac1 constructs, Q61L and Q61L/G12V, exhibit markedly stronger interaction with csRAPH of Lpds than wild-type Rac1(**Fig. 2D**). These results confirm that Rac1 directly interacts with the csRAPH region of Lpds in a GTPase activity-dependent manner.

### The Lpds:Rac1 complex adopts a non–β-sheet binding mode distinct from the Rap1:RIAM interaction

We then probed the binding mode of Rac1 with the csRAPH region of Lpds. Since crystallization attempts of the Rac1:csRAPH complex or their fusion protein were unsuccessful, we performed structure prediction of csRAPH using Alpha-Fold.^36^ The predicted structure was then aligned with the RAPH structure of RIAM from the RIAM:RAP1 complex crystal structure (PDB ID: 4KVG).^31^ In the predicted model, the CC region and the RA-PH module are consistent with the previously reported crystal structure of the CC-RA-PH fragment of the main Lpd isoform.^4^ The N-terminal half of the CS insertion is largely unstructured, while the C-terminal half of the CS forms two short α-helices that contact with CC region, likely stabilizing its helical configuration (**Fig. 3A**). The RA-PH module of Lpds closely resembles that of RIAM in the RIAM:Rap1 complex, with an RMSD of 0.7-Å over all atoms.

**Figure 3.**
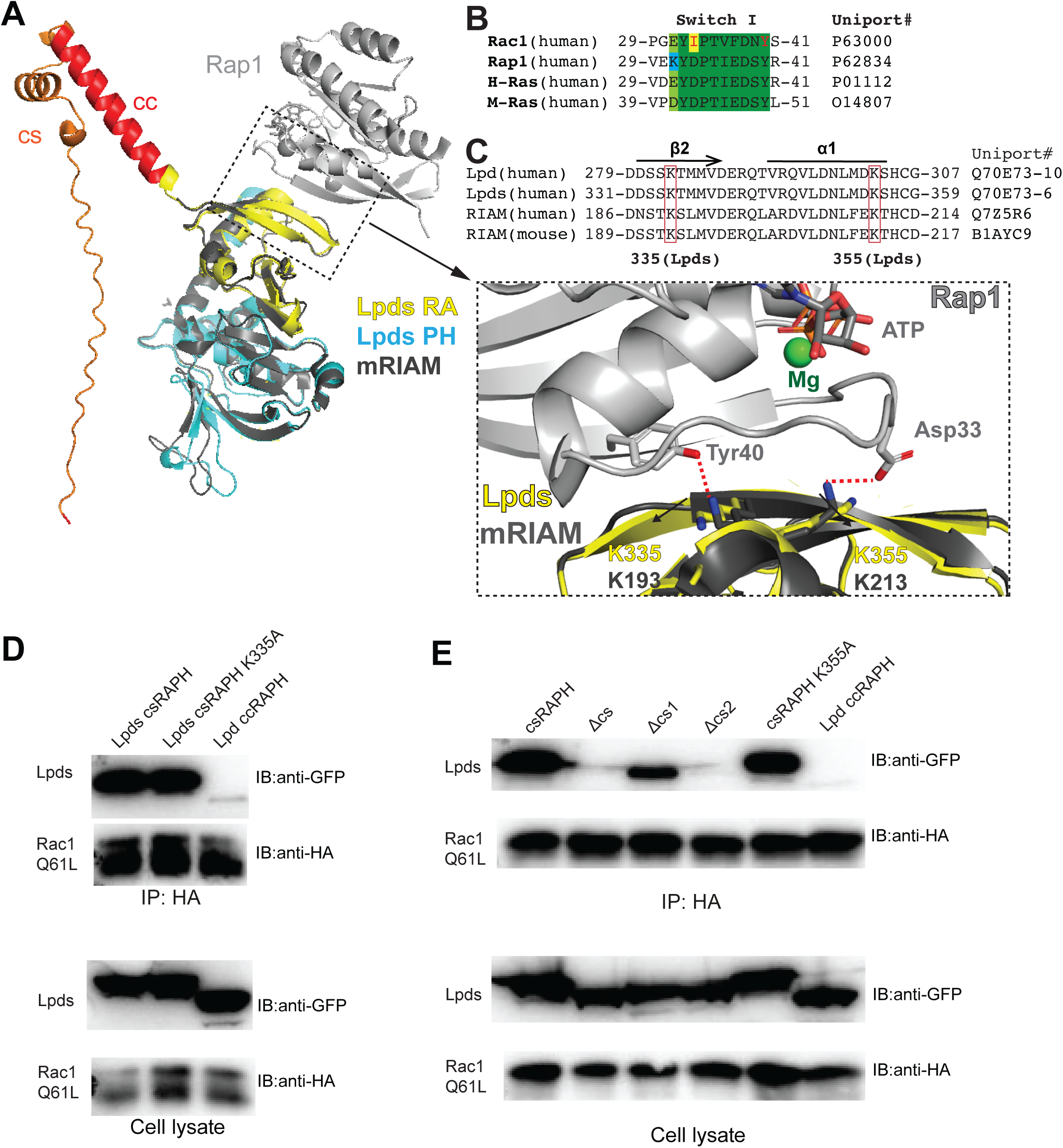
Lpds binds Rac1 through a non–β-sheet interface distinct from the RIAM:Rap1 interaction. (**A**). Structural alignment of the AlphaFold-predicted Lpds model with the crystal structure of the RIAM:Rap1 complex. Lpds domains are colored as follows: CS insertion (orange), CC (red), RA domain (yellow), and PH domain (cyan). The RIAM:Rap1 complex is shown in gray. The zoomed-in panel (bottom right) highlights key salt bridges in the RIAM:Rap1 interface (Asp33:Lys213 and Tyr40:Lys193). The corresponding residues in Lpds (Lys335 and Lys355), selected for mutagenesis, are labeled. (**B**). Sequence alignment of the Switch I region from Rac1, Rap1, H-Ras, and M-Ras. Conserved residues are highlighted in green. Rac1 residues Ile33 and Tyr40, corresponding to salt bridge–forming residues in Rap1, are marked in red. (**C**) Sequence alignment of Lpd, Lpds and RIAM, with lysine residues involved in Rap1:RIAM interaction highlighted in the boxes. Lpds numbering is based on the sequence includes the CS insertion. (**D**). Co-IP assay of HA-tagged Rac1 Q61L with GFP-tagged Lpds csRAPH, csRAPH K335A, and Lpd ccRAPH. (**E**). Co-IP assay of HA-tagged Rac1 Q61L with various truncations of GFP-tagged Lpds csRAPH and csRAPH K355A.

Interestingly, structural modeling suggests that if the Rac1:Lpds interaction resembles a canonical anti-parallel β-sheet interaction of typical RA:GTPase interactions as observed in the Rap1:RIAM complex, no direct contact involving the CS region would be expected (**Fig. 3A**). In the Rap1:RIAM complex structure, two hydrogen-bond interactions are observed between RIAM-Lys193 and Rap1-Tyr40, and between RIAM-Lys213:Rap1-Asp33. While both Asp33 and Tyr40 are highly conserved across Ras family GTPases, only Tyr40 remains conserved in the Rho family (**Fig. 3B**). Although the corresponding lysine residues are conserved in Lpds as Lys335 and Lys355 (residue numbering including the cs insertion) (**Fig. 3C**), substitution of Asp33 by isoleucine in Rac1 prevents the formation of the signature Asp-Lys hydrogen bond,^31^ suggesting that the Rac1:Lpds interaction adopts a noncanonical RA:Ras binding mode. To test this, we mutated Lys355 in Lpds, which corresponds to RIAM-Lys213 in RIAM that forms the hydrogen bond with Rap1-Asp33 in the Rap1:RIAM complex. Lpds K355A exhibits similar binding affinity to Rac1 as WT-Lpds, supporting the absence of the signature Asp-Lys or any compensatory hydrogen bond in the Rac1:Lpds interaction (**Fig. 3D**). Moreover, we generated a K335A mutation, which also had minimal effect on Rac1:Lpds interaction, further supporting that the Rac1:Lpds interaction adopts a noncanonical RA:Ras binding mode distinct from the RIAM:Rap1 interaction. (**Fig. 3E**)

### Leucine Clusters in the CS region are critical for the non-canonical Rap1:Lpds interactions

The CS insertion in Lpds includes two alternative splicing segments, cs1 and cs2. Depending on the splicing product, each isoform may contain cs1 alone (isoforms #3, #7), cs2 alone (isoforms #4, #8), both cs1 and cs2 (CS, isoforms #6, #9), or neither (isoforms #2, #5, #10) (**Fig. 1A**). Although secondary structure prediction suggests a weak helical tendency in cs1, Alphafold prediction indicates a largely unstructured cs1 region (**Fig. 3A**). In contrast, cs2 is predicted to form two short α-helices and contains two leucine clusters, Leu10’-Leu11’-Leu12’ (3L) and Leu22’-Leu23’ (2L) (residue numbering within the cs2 insertion only), within the two helices (**Fig. 4A**). To test the impact of cs1 and cs2 on Rac1 binding, we generated a series of deletion constructs derived from the csRAPH segment by removing either cs1 (ι1cs1) or cs2 (ι1cs2), or both (ι1CS), and performed co-IP assays with constitutively active Rac1 (**Fig. 3E**). Removal of cs1 had minimal impact on Rac1 binding, whereas removal or cs2 or removal of both cs1 and cs2 completely abolished the interaction between Lpds and Rac1, indicating that cs2 plays an essential role in mediating Rac1 binding.

**Figure 4.**
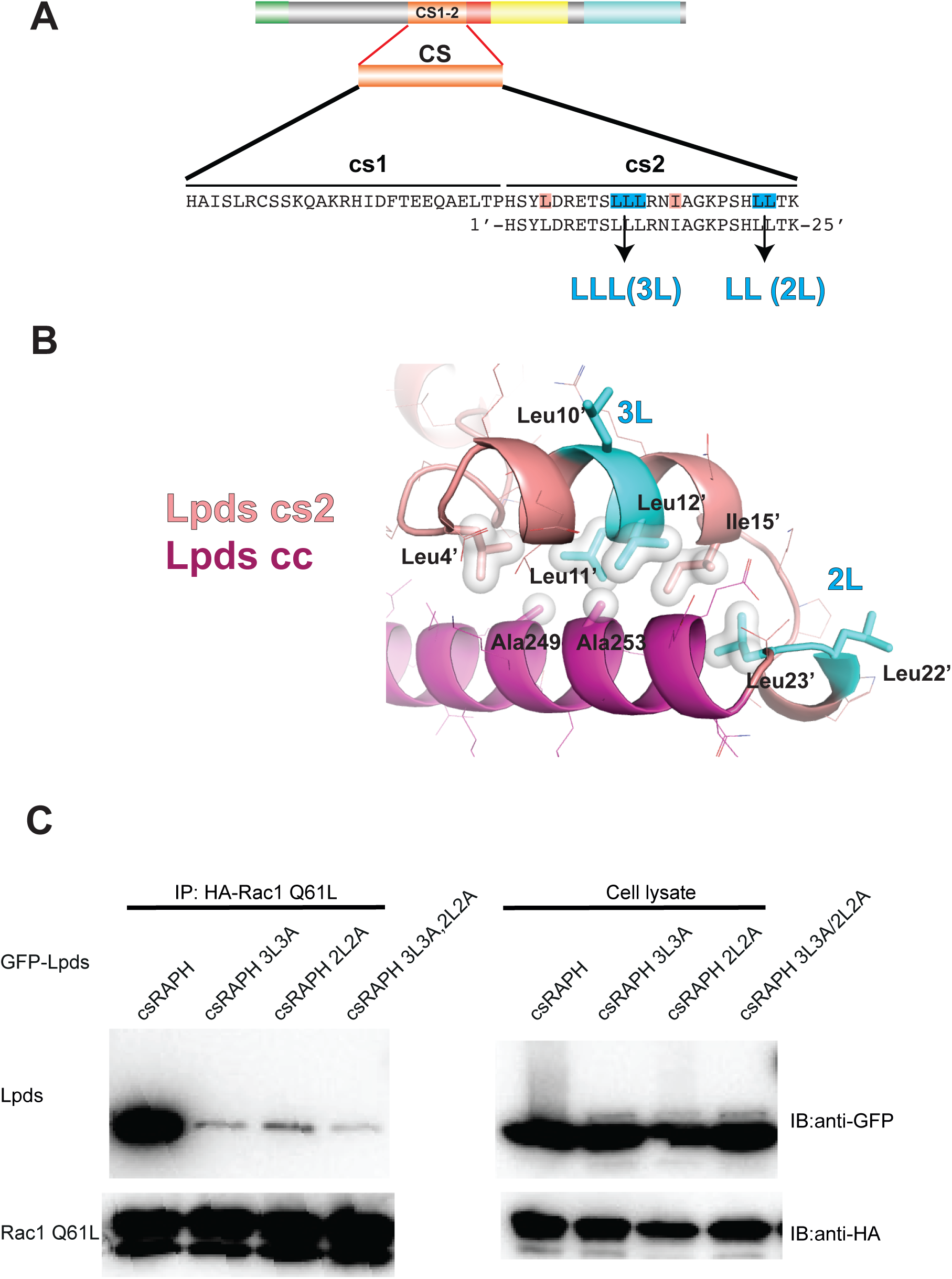
Structure prediction and co-IP assay suggest that the cs2 insertion stabilize the CC region via hydrophobic interaction. (A). Amino acid sequence of the two Lpd splicing insertions, cs1 and cs2, are shown. The cs2 insertion residues are numbered 1’ to 25’. Two hydrophobic leucine clusters, LLL (3L) and LL (2L), in the cs2 insertion are highlighted in cyan. Leu4’ and Ile15’ of cs2 are colored in pink. (**B**). Predicted interface between the cs2 insertion (in pink) and CC region (in red). Hydrophobic residues contributing to the cs2:CC interaction are shown in surface representation. Residue numbers of CC are based on the Lpd main isoform to avoid confusion. (**C**). Co-IP assay of HA-tagged Rac1 Q61L and GFP-tagged Lpds csRAPH, Lpds csRAPH 3L3A, Lpds csRAPH 2L2A and Lpds csRAPH 3L3A/2L2A.

Leucine clusters are often involved in hydrophobic interaction and helix stabilization. We then investigated the role of these leucine clusters in Rac1 binding. In the predicted model, the 3L and 2L are positioned near Leu4’ and Ile15’ in the cs2 and Ala 249and Ala253 in the CC region (**Fig. 4B**), suggesting that they help stabilize the CC helix via intramolecular hydrophobic packing. To test the impact of the Leu clusters, we generated 3L3A (LLL to AAA) and 2L2A (LL to AA) mutations. Co-IP assays using the constitutively active Rac1-Q61L demonstrate that the disruption of either clusters or both significantly diminishes Rac1 binding (**Fig. 4C**). These results indicate that both leucine clusters contribute to either direct interaction with Rac1 or stabilizing the structural configuration required for the interaction

### Structure prediction reveals a noncanonical Rac1:Lpds interaction mediated by a cs2-stabilized helical surface

Although we were unable to obtain a crystal structure of the Lpds:Rac1 complex, we performed Alphafold3 structure predictions using multiple Lpds constructs and GTP-bound Rac1. All predicted models consistently reveal a conserved interface between the cs2-CC region of Lpds and the active form of Rac1, in strong agreement with our biochemical data. These findings suggest that the predicted interaction of the Lpds:Rac1 complex is structurally robust and biologically reliable.

The predicted Lpds:Rac1 complex structure reveals a strikingly unique binding mode of GTPase:effector. Most of small GTPases in the Rho and Ras family engage their effector proteins via a canonical β-sheet interaction with the RA of RBD (Ras-binding domain) or a helical-pair interaction. In the classic helical-pair interaction such as the Rac1:Prk1 complex (PDB ID: 2RMK, **Fig. 5A)**, a pair of α-helices from the effector contact the Switch I and Switch II regions of the GTPase. The helical pairs are stabilized through coiled-coil interaction or other interhelical packing, forming an extensive interface with the target GTPase. In contrast, although all Lpd isoforms contain a CC region that mediates an intermolecular coiled-coil in a homo-dimer formation,^4^ no significant interaction is observed between Lpd main isoform and Rac1, indicating that the Lpds:Rac1 interaction is not mediated by the canonical helical-pair binding mode. Instead, structure prediction suggests that the CC region of Lpds is stabilized by the cs2 insertion, forming a hydrophobic surface that directly interacts with the corresponding hydrophobic surface between Switch I and Switch II regions in the active Rac1(**Fig. 5B**). The interaction is mediated by residues Val36, Phe37, Leu67, and Leu70 in the Switch regions and residue Trp56 from Rac1 and residues Leu256, Leu260, Ile263 in CC (**Fig. 5B**). This interface is further reinforced by Tyr3’ and Leu4’ residues from the cs2 insertion. Thus, unlike the classical helical-pair interaction, Lpds:Racl interaction reveals a noncanonical, splicing insertion-dependent, single-helix binding mode that is not previously observed.

**Figure 5.**
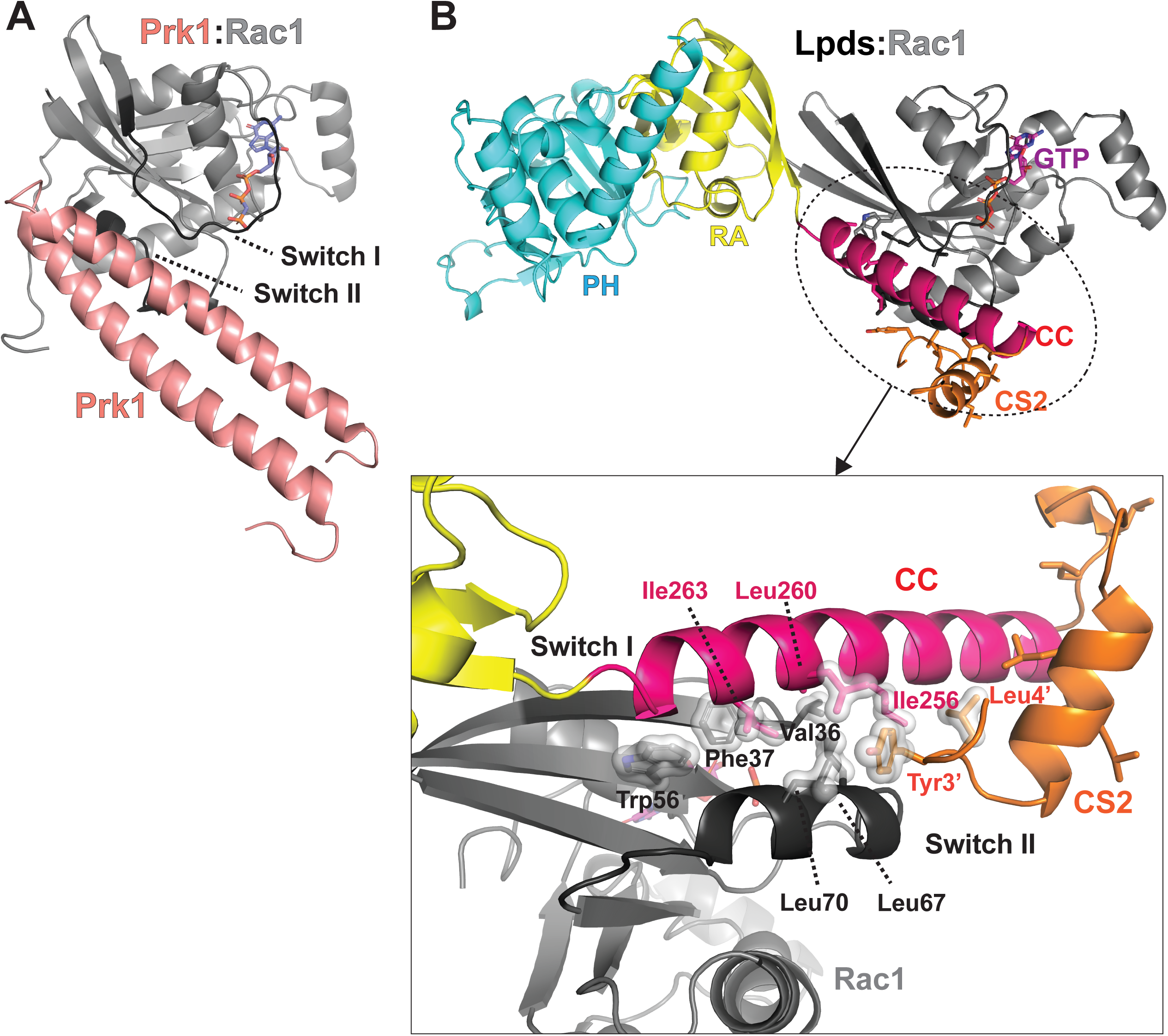
AlphaFold3–predicted Lpds:Rac1 structure reveals a noncanonical single-helix binding mode. (B). Structure of Rac1 (in grey) in complex with the helical-pair for Prk1 (in pink). (**B**). Structure Lpds:Rac1 complex predicted by AlphaFold3. *Upper*: Lpds is colored as indicated in Fig. 1A. Rac1 is colored in Gray. The interface between Lpds-cs2-CC and Rac1 is highlighted in a zoomed-in view below. *Lower*: Switch I and Switch II regions of Rac1 are colored in dark grey. Residues forming the hydrophobic interaction interface are shown in surface representation. These includes Ile256, Leu260, and Ile263 (in red) from the Lpds CC region; residues 3 (Tyr3’) and 4 (Leu4’) from the cs2 insertion (in orange); Val36 and Phe37 from Switch I of Rac1 (in grey); and Leu67 and Leu70 from Switch II of Rac1 (in dark grey); and Trp56 in Rac1.

## Discussion

Small GTPases are ubiquitously expressed across tissues and play central roles in regulating cell growth, differentiation, and migration. Among them, Ras family members such as H-Ras, N-Ras, and K-Ras are well-established oncogenes, frequently mutated in cancer to promote uncontrolled proliferation through persistent GTP-bound activation. In contrast, Rho family GTPases, including Rac1, are primarily associated with cancer metastasis due to their regulation of cytoskeletal dynamics. Multiple cancer-associated mutations in Rac1 have been identified, including the melanoma hotspot mutation P29S.^37^ Rac1-P29S is shown to increase the flexibility of Switch I.^38^ P29S can activate Rac1 by substantially increasing inherent GDP/GTP nucleotide exchange, resulting in fast cycling of GTPase.^39^ These changes enhance Rac1 binding with downstream effectors and promoted melanocyte proliferation and migration.^40^ Moreover, the P29S mutation is also involved in melanoma resistance to RAF inhibitors,^41^ and may contribute to antitumor immune response by altering the expression level of death-ligand 1(PD-L1).^37^ Another mutation, A159V, has been shown to be associated with head and neck squamous cell carcinoma (HNSCC).^42^ ^43^ Overall, Rac1 is emerging as both hotspot mutations and as a promising therapeutic biomarker in selected cancers.

As a Rac1 GTPase effector, Lpd can recruit Ena/VASP proteins to the leading edge of lamellipodia and filopodia, thus regulating cell migration by controlling actin polymerization and dissociation.^19,20^ Lpd, and its mammalian ortholog RIAM, possess a key structural module comprising an RA domain, a PH domain, and an upstream coiled-coil region. Through its interation with Rac1, Lpd modulates Scar/WAVE complex and regulates cell migration.^19^ We find that the cs2 insertion in the Lpds isoform is required for Rac1 binding likely by stablizing a hydrophic interface between the CC region and Rac1.

This cs2 insertion is absent in the Lpd main isoform. Conversely,the Ena/VASP-binding region in the Lpd main isoform is also missing in the Lpds isoform. Based on our findings, we propose that Lpd contributes to Rac1-mediated actin regulation through two complementary mechanisms: Lpds engages Rac1 signaling via the cs2-enhanced single-helix binding mode, while the Lpd main isoform regulates actin polymerization machinery via Ena/VASP recruitment. These distinct functions may be coordinated through Lpd dimerization or by interactions with other actin-regulatory complexes.

Previous studies have revealed expression of Lpds and Lpd in fibroblast.^19^ To investigate the role Lpd in cancer, we analyzed the expression levels of Lpd isofroms in various cancer cells (**Supplemental Figure**). We found that the short isoform Lpds shows much higher expression in several cancer types, such as HeLa (cervical cancer), H446 (lung cancer), and PNX001 and PNX017 (pancreatic cancer). In contrast, its expression is very low or not detectable in normal tissues, including adult brain and fibroblast-derived lines like MR90 and RPE1. The main Lpd isoform exhibits more consistent expression, especially in non-cancerous cells. This result suggests that cancer cells may selectively express Lpds, which contains a special cs2 insertion that enhances Rac1 binding in a GTPase activity-dependent manner. Nevertheless, Lpds lacks the Ena/VASP-binding region present in the main Lpd isoform, indicating that these two isoforms may regulate actin and cell migration through different mechanisms. This isoform-specific expression pattern may have important implications for understanding cancer behavior. Lpds may serve as a biomarker for invasive or metastatic cancer, and also as a potential target for therapy. It will be important to examine whether Lpds supports tumor-specific Rac1 signaling, and whether blocking this isoform can reduce cancer cell migration without affecting normal cells. Together, our findings provide new insight into how Rac1 signaling is regulated in cancer, and suggest a possible direction for developing isoform-targeted therapies.

## Method

### Plasmid construction

Lamellipodin constructions were subcloned into a modified pET28a vector with a hexahistidine-tag (His6-tag). The Rac1s were subcloned into pGEX-5X-1 vector with a GST-tag. Lamellipodin was subcloned into an EFEP-C3 vector and Rac1 was subcloned into pCGN vector for expression in HEK293 cells. Point mutations were generated with a site-directed mutagenesis according to the QuikChange Site-directed mutagenesis manual.

### Protein purification

The His6-tagged proteins and GST-tagged proteins were expressed and purified according to the procedures described previously. (Gao, cell reports, Gao structure, Zhang PNAS) Briefly, *E. coli* BL21(DE3) was transformed with pET28a-Lpd constructs or pGEX-Rac1 constructs. The positive colonies were cultured in LB medium with 50µg/ml kanamycin or 80µg/ml ampicillin in shaking flasks (200 rpm/min) at 37℃. To induce protein expression, 0.2mM isopropyl-D-1-thiogalactopyranoside (IPTG) was added to the flasks when the OD600 reached 0.7. For His6-tagged protein production, the flasks kept shaking overnight at 16℃ and for GST-tagged protein production, the flasks kept shaking at 37℃ for 3H. The bacteria were collected by spinning down at 2200rpm/min for 30 min and were resuspended with 20mM Tris pH 7.5, 500mM NaCl for His6-tagged proteins and 20mM Tris pH 7.5, 100mM NaCl for GST-tagged proteins. EmulsiFlex-C3 was used to homogenize the resuspension, and the supernatant were applied to HisTrap FF columns (Cytiva) or GSTrap HP columns (Cytiva) for purification using ÄKTA Purifier (Cytiva).

### Co-immunoprecipitation

GFP-tagged Lpd proteins and HA-tagged Rac1 proteins were co-expressed in HEK293 cells. The cell lysates were rotating with α-HA (sigma, H9658-.2ML)-conjugated Surebeads^TM^ magnetic beads (161-4023, Bio-Rad, Hercules, CA, USA) for 2H at room temperature. The supernatant was removed, and the beads were washed with PBS-T (PBS+0.1% Tween 20) three times. Samples were eluted by incubating the beads with glycine (20mM, pH 2.0) for 5min at room temperature. After mixing with Laemmli sample buffer (S3401-10VL) and heated to 95℃ for 10min, the samples were applied to Western Blotting. Anti-GFP (Sigma, G1544, 1:2000) and anti-HA (sigma, H9658-.2ML, 1:2000) were used for the detection of GFP-tagged Lpd proteins and HA-tagged Rac1 proteins. A FluorChem E System (Proteinsimple) with a charge-coupled device (CCD) camera was used to expose the Western blotting membrane.

### GST pull-down

Purified GST-Rac1 proteins (100ul, 1mg/ml) were incubated with 2mM EDTA on ice for 30min then incubated with 5mM GTP and 20mM MgCl2 on ice for 30min. His6-Lpd proteins (100ul, 1mg/ml) were mixed with GTP-loaded GST-Rac1 proteins in binding buffer (50 mM Tris, pH 7.5, 100 mM NaCl, 2 mM DTT). The protein mixture was rotating for 1 hour at 4℃ then incubated with binding buffer-equilibrated glutathione agarose beads (Invitrogen, G2879) on a rotator for 1 hour at 4 ℃. The beads were washed with binding buffer three times after the supernatant is removed. The beads-binding proteins were eluted with 40ul elution buffer (50 mM Tris, pH 7.5, 100 mM NaCl, 2 mM DTT, 10 mM reduced glutathione). Coomassie staining and Western Blotting were used to evaluate the result after samples were applied to SDS/PAGE. Anti-His (Sigma, H1029-100UL, 1:2000) was used for the detection of His6-tagged THD proteins.

## Acknowledgments

We thank Dr. Zeng-jie Yang (Fox Chase Cancer Center) for providing the tumor cell samples and the discussions. We thank the beamline staff of AMX and FMX at National Synchrotron Light Source-II, Brookhaven National Laboratory for technical support. This work was supported by an NIH Grant GM119560 (to J.W.), an ASH bridge grant (to J.W.), a Pennsylvania Department of Health Grant 4100085739 (to J.W.), ACS RSG-15-167-01-DMC (to J.W.). T.G. was partially supported by the Elizabeth Knight Patterson Postdoctoral Fellowship.

## Author Contributions

T.G. and P.Z. carried out protein production and the biochemical assays. T.G. performed structural analyses, expression tests, and mutagenesis. A.M.K and J.S.D performed mass spectrometry. T.G. and J.W. wrote the manuscript and are the co-corresponding authors. J.W. conceived and supervised the project.

---

**Supplymental Figure.**
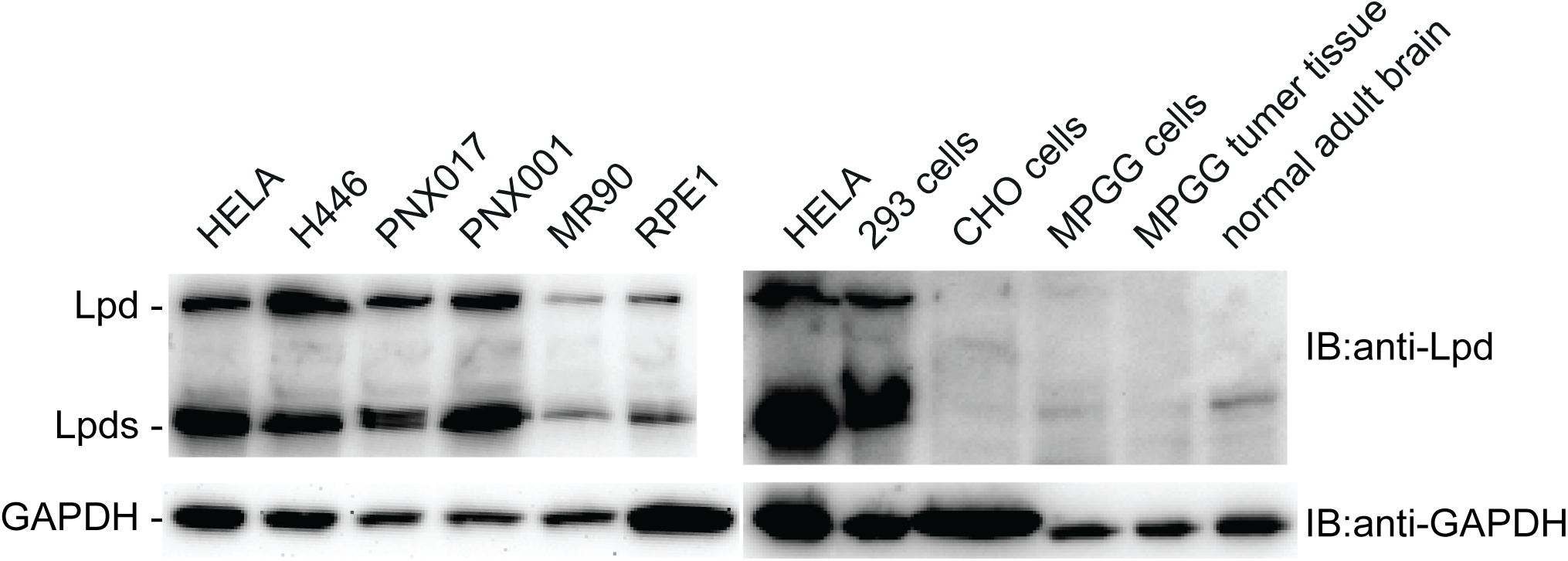
Expression level of isoforms of Lpd in different cells lines. Lpd isoforms expression in HeLa (cervical cancer), H446(lung cancer), PNX001(pancreatic cancer), PNX017(pancreatic cancer), IMR90(fibrocyte), RPE1(fibrocyte), 293 (kidney cell),CHO (Chinese hamster ovary cell), normal brain tissue and MPGG tumor tissue.

## References

1. Hansen, S.D., and Mullins, R.D. (2015). Lamellipodin promotes actin assembly by clustering Ena/VASP proteins and tethering them to actin filaments. Elife 4. 10.7554/eLife.06585.

2. Carmona, G., Perera, U., Gillett, C., Naba, A., Law, A.L., Sharma, V.P., Wang, J., Wyckoff, J., Balsamo, M., Mosis, F., et al. (2016). Lamellipodin promotes invasive 3D cancer cell migration via regulated interactions with Ena/VASP and SCAR/WAVE. Oncogene 35, 5155–5169. 10.1038/onc.2016.47.

3. Sari-Ak, D., Torres-Gomez, A., Yazicioglu, Y.F., Christofides, A., Patsoukis, N., Lafuente, E.M., and Boussiotis, V.A. (2022). Structural, biochemical, and functional properties of the Rap1-Interacting Adaptor Molecule (RIAM). Biomed J 45, 289–298. 10.1016/j.bj.2021.09.005.

4. Chang, Y.C., Zhang, H., Brennan, M.L., and Wu, J. (2013). Crystal structure of Lamellipodin implicates diverse functions in actin polymerization and Ras signaling. Protein Cell 4, 211–219. 10.1007/s13238-013-2082-5.

5. Chang, Y.C., Zhang, H., Franco-Barraza, J., Brennan, M.L., Patel, T., Cukierman, E., and Wu, J. (2014). Structural and mechanistic insights into the recruitment of talin by RIAM in integrin signaling. Structure 22, 1810–1820. 10.1016/j.str.2014.09.020.

6. Chang, Y.-C., Su, W., Cho, E.-a., Zhang, H., Huang, Q., Philips, M.R., and Wu, J. (2019). Molecular basis for autoinhibition of RIAM regulated by FAK in integrin activation. Proceedings of the National Academy of Sciences 116, 3524–3529.

7. Lyulcheva, E., Taylor, E., Michael, M., Vehlow, A., Tan, S., Fletcher, A., Krause, M., and Bennett, D. (2008). Drosophila pico and its mammalian ortholog lamellipodin activate serum response factor and promote cell proliferation. Dev Cell 15, 680–690. 10.1016/j.devcel.2008.09.020 S1534-5807(08)00402-4 [pii].

8. Gray, J.L., von Delft, F., and Brennan, P.E. (2020). Targeting the Small GTPase Superfamily through Their Regulatory Proteins. Angew Chem Int Ed Engl 59, 6342–6366. 10.1002/anie.201900585.

9. Etienne-Manneville, S., and Hall, A. (2002). Rho GTPases in cell biology. Nature 420, 629–635.

10. Perez, P., and Rincon, S.A. (2010). Rho GTPases: regulation of cell polarity and growth in yeasts. Biochem J 426, 243–253. 10.1042/BJ20091823.

11. Sadok, A., and Marshall, C.J. (2014). Rho GTPases: masters of cell migration. Small GTPases 5, e29710. 10.4161/sgtp.29710.

12. Villalonga, P., and Ridley, A.J. (2006). Rho GTPases and cell cycle control. Growth Factors 24, 159–164. 10.1080/08977190600560651.

13. De, P., Aske, J.C., and Dey, N. (2019). RAC1 Takes the Lead in Solid Tumors. Cells 8. 10.3390/cells8050382.

14. Kotelevets, L., and Chastre, E. (2020). Rac1 signaling: from intestinal homeostasis to colorectal cancer metastasis. Cancers 12, 665.

15. Seiz, J.R., Klinke, J., Scharlibbe, L., Lohfink, D., Heipel, M., Ungefroren, H., Giehl, K., and Menke, A. (2020). Different signaling and functionality of Rac1 and Rac1b in the progression of lung adenocarcinoma. Biol Chem 401, 517–531. 10.1515/hsz-2019-0329.

16. Law, A.-L., Vehlow, A., Kotini, M., Dodgson, L., Soong, D., Theveneau, E., Bodo, C., Taylor, E., Navarro, C., and Perera, U.J.J.C.B. (2013). Lamellipodin and the Scar/WAVE complex cooperate to promote cell migration in vivo. 203, 673–689.

17. Tasaka, G.-i., Negishi, M., and Oinuma, I.J.J.o.N. (2012). Semaphorin 4D/Plexin-B1-mediated M-Ras GAP activity regulates actin-based dendrite remodeling through Lamellipodin. 32, 8293–8305.

18. Tasaka, G., Negishi, M., and Oinuma, I. (2012). Semaphorin 4D/Plexin-B1-mediated M-Ras GAP activity regulates actin-based dendrite remodeling through Lamellipodin. J Neurosci 32, 8293–8305. 10.1523/JNEUROSCI.0799-12.2012.

19. Krause, M., Leslie, J.D., Stewart, M., Lafuente, E.M., Valderrama, F., Jagannathan, R., Strasser, G.A., Rubinson, D.A., Liu, H., Way, M., et al. (2004). Lamellipodin, an Ena/VASP ligand, is implicated in the regulation of lamellipodial dynamics. Dev Cell 7, 571–583. 10.1016/j.devcel.2004.07.024.

20. Dimchev, G., Amiri, B., Humphries, A.C., Schaks, M., Dimchev, V., Stradal, T.E.B., Faix, J., Krause, M., Way, M., Falcke, M., and Rottner, K. (2020). Lamellipodin tunes cell migration by stabilizing protrusions and promoting adhesion formation. J Cell Sci 133. 10.1242/jcs.239020.

21. Law, A.L., Vehlow, A., Kotini, M., Dodgson, L., Soong, D., Theveneau, E., Bodo, C., Taylor, E., Navarro, C., Perera, U., et al. (2013). Lamellipodin and the Scar/WAVE complex cooperate to promote cell migration in vivo. J Cell Biol 203, 673–689. 10.1083/jcb.201304051/jcb.201304051 jcb.201304051 [pii].

22. Moritz, S., Krause, M., Schlatter, J., Cordes, N., and Vehlow, A. (2021). Lamellipodin-RICTOR Signaling Mediates Glioblastoma Cell Invasion and Radiosensitivity Downstream of EGFR. Cancers (Basel) 13. 10.3390/cancers13215337.

23. Yasui, H., Ohnishi, Y., Kakudo, K., NOZAKI, M., and Nakajima, M. (2017). HGF/cMet induces cell migration of oral squamous cell carcinoma via lamellipodin. Journal of Osaka Dental University 51, 1–8.

24. Eppert, K., Wunder, J.S., Aneliunas, V., Tsui, L.C., Scherer, S.W., and Andrulis, I.L. (2005). Altered expression and deletion of RMO1 in osteosarcoma. Int J Cancer 114, 738–746. 10.1002/ijc.20786.

25. Quinn, C.C., Pfeil, D.S., and Wadsworth, W.G. (2008). CED-10/Rac1 mediates axon guidance by regulating the asymmetric distribution of MIG-10/lamellipodin. Curr Biol 18, 808–813. 10.1016/j.cub.2008.04.050 S0960-9822(08)00536-8 [pii].

26. Mott, H.R., and Owen, D. (2015). Structures of Ras superfamily effector complexes: What have we learnt in two decades? Crit Rev Biochem Mol Biol 50, 85–133. 10.3109/10409238.2014.999191.

27. Stastna, M., and Van Eyk, J.E. (2012). Analysis of protein isoforms: can we do it better? Proteomics 12, 2937–2948. 10.1002/pmic.201200161.

28. Chang, Y.C., Zhang, H., Franco-Barraza, J., Brennan, M.L., Patel, T., Cukierman, E., and Wu, J. (2014). Structural and Mechanistic Insights into the Recruitment of Talin by RIAM in Integrin Signaling. Structure E-published.

29. Chang, Y.C., Su, W., Cho, E.A., Zhang, H., Huang, Q., Philips, M.R., and Wu, J. (2019). Molecular basis for autoinhibition of RIAM regulated by FAK in integrin activation. Proc Natl Acad Sci U S A 116, 3524–3529. 10.1073/pnas.1818880116.

30. Cheng, K.W., and Mullins, R.D. (2020). Initiation and disassembly of filopodia tip complexes containing VASP and lamellipodin. Mol Biol Cell 31, 2021–2034. 10.1091/mbc.E20-04-0270.

31. Zhang, H., Chang, Y.C., Brennan, M.L., and Wu, J. (2014). The structure of Rap1 in complex with RIAM reveals specificity determinants and recruitment mechanism. J Mol Cell Biol 6, 128–139. 10.1093/jmcb/mjt044 mjt044 [pii].

32. Wynne, J.P., Wu, J., Su, W., Mor, A., Patsoukis, N., Boussiotis, V.A., Hubbard, S.R., and Philips, M.R. (2012). Rap1-interacting adapter molecule (RIAM) associates with the plasma membrane via a proximity detector. J Cell Biol 199, 317–330. 10.1083/jcb.201201157 jcb.201201157 [pii].

33. Depetris, R.S., Wu, J., and Hubbard, S.R. (2009). Structural and functional studies of the Ras-associating and pleckstrin-homology domains of Grb10 and Grb14. Nat Struct Mol Biol 16, 833–839. 10.1038/nsmb.1642 nsmb.1642 [pii].

34. Colicelli, J. (2004). Human RAS Superfamily Proteins and Related GTPases. Science’s STKE 2004. 10.1126/stke.2502004re13.

35. Hood, F.E., Sahraoui, Y.M., Jenkins, R.E., and Prior, I.A. (2023). Ras protein abundance correlates with Ras isoform mutation patterns in cancer. Oncogene 42, 1224–1232.

36. Jumper, J., Evans, R., Pritzel, A., Green, T., Figurnov, M., Ronneberger, O., Tunyasuvunakool, K., Bates, R., Zidek, A., Potapenko, A., et al. (2021). Highly accurate protein structure prediction with AlphaFold. Nature 596, 583–589. 10.1038/s41586-021-03819-2.

37. Vu, H.L., Rosenbaum, S., Purwin, T.J., Davies, M.A., and Aplin, A.E. (2015). RAC1 P29S regulates PD-L1 expression in melanoma. Pigment Cell Melanoma Res 28, 590–598. 10.1111/pcmr.12392.

38. Senyuz, S., Jang, H., Nussinov, R., Keskin, O., and Gursoy, A. (2021). Mechanistic Differences of Activation of Rac1(P29S) and Rac1(A159V). J Phys Chem B 125, 3790–3802. 10.1021/acs.jpcb.1c00883.

39. Davis, M.J., Ha, B.H., Holman, E.C., Halaban, R., Schlessinger, J., and Boggon, T.J. (2013). RAC1P29S is a spontaneously activating cancer-associated GTPase. Proceedings of the National Academy of Sciences 110, 912–917.

40. Krauthammer, M., Kong, Y., Ha, B.H., Evans, P., Bacchiocchi, A., McCusker, J.P., Cheng, E., Davis, M.J., Goh, G., Choi, M., et al. (2012). Exome sequencing identifies recurrent somatic RAC1 mutations in melanoma. Nat Genet 44, 1006–1014. 10.1038/ng.2359.

41. Watson, I.R., Li, L., Cabeceiras, P.K., Mahdavi, M., Gutschner, T., Genovese, G., Wang, G., Fang, Z., Tepper, J.M., Stemke-Hale, K., et al. (2014). The RAC1 P29S hotspot mutation in melanoma confers resistance to pharmacological inhibition of RAF. Cancer Res 74, 4845–4852. 10.1158/0008-5472.CAN-14-1232-T.

42. Ngan, H.L., Liu, Y., Poon, P.H.Y., and Lui, V.W.Y. (2019). RAC1 genomic aberrations as predictive biomarkers for head and neck squamous cell carcinoma (HNSCC). Cancer Research 79, 4033–4033.

43. Chang, M.T., Asthana, S., Gao, S.P., Lee, B.H., Chapman, J.S., Kandoth, C., Gao, J., Socci, N.D., Solit, D.B., Olshen, A.B., et al. (2016). Identifying recurrent mutations in cancer reveals widespread lineage diversity and mutational specificity. Nat Biotechnol 34, 155–163. 10.1038/nbt.3391.

